# DEBay: a computational tool for deconvolution of quantitative PCR data for estimation of cell type-specifc gene expression in a mixed population

**DOI:** 10.1101/2020.04.10.035642

**Authors:** Vimalathithan Devaraj, Biplab Bose

## Abstract

The expression of a gene is commonly estimated by quantitative PCR (qPCR) using RNA isolated from a large number of pooled cells. Such pooled samples often have subpopulations of cells with different levels of expression of the target gene. Estimation of gene expression from an ensemble of cells obscures the pattern of expression in different subpopulations. Physical separation of various subpopulations is a demanding task. We have developed a computational tool, Deconvolution of Ensemble through Bayes-approach (DEBay), to estimate cell type-specific gene expression from qPCR data of a mixed population. DEBay estimates Normalized Gene Expression Coefficient (NGEC), which is a relative measure of the expression of the target gene in each cell type in a population. NGEC has a direct algebraic correspondence with the normalized fold change in gene expression measured by qPCR. DEBay can deconvolute both time-dependent and -independent gene expression profiles. It uses the Bayesian method of model selection and parameter estimation. We have evaluated DEBay using synthetic and real experimental data. DEBay is implemented in Python. A GUI of DEBay and its source code are available for download at SourceForge (https://sourceforge.net/projects/debay).

## 1. Introduction

A population of cells often contain several subpopulations with distinct gene expression pattern. However, gene expression experiments using pooled cells, like quantitative PCR (qPCR), expression microarray, obscure the information on subpopulation- or cell-type-specific gene expression. For example, in an *in vitro* experiment on epithelial-to-mesenchymal transition, one may have samples having mixtures of epithelial and mesenchymal cells. Each of these cell types has a distinct gene expression signature. However, a qPCR experiment would not be able to provide such cell type-specific information on gene expression. A similar problem is faced in the analysis of gene expression in biopsy samples of solid tumors. A tumor sample is usually heterogeneous with different subpopulations (Hinohara and Polyak, 2019; Marusyk and Polyak, 2010). However, qPCR experiments using bulk tumor samples would provide us the average gene expression of all these cells. Further, tumor samples often contain tumor-infiltrating lymphocytes and endothelial cells (Hida, et al., 2018; Pages, et al., 2010). These infiltrating cells confound the gene expression analysis of cancer cells when the bulk tumor is used as a sample (de Ridder, et al., 2005; Kuhn, et al., 2011).

This problem can be alleviated if we physically segregate different subpopulations and investigate the gene expression in each of them. Physical separation techniques have limitations like poor yield and quality of RNA (Debey, et al., 2004; Zhong, et al., 2013). Separation of different subpopulations is difficult when subpopulations are defined in terms of physical properties like-morphology, motility, or functions (Devaraj and Bose, 2019; Kimmel, et al., 2018). Computational algorithms to deconvolute population-level data into subpopulation-specific information help in these circumstances.

A gene expression deconvolution algorithm requires two inputs: a) population-level gene expression data, and b) the proportions or relative size of each subpopulation in the sample. Note that in certain experiments relative size of each subpopulation in a mixture of cells can be estimated even without physical segregation of these cells (Devaraj and Bose, 2019; Kimmel, et al., 2018; Mandal, et al., 2016). With this information, a deconvolution algorithm estimates relative gene expression in each subpopulation in a heterogeneous sample.

Several authors have used the deconvolution approach to estimate the cell type-specific gene expression patterns from microarray data (Dimitrakopoulou, et al., 2018; Erkkila, et al., 2010; Lahdesmaki, et al., 2005; Zhong, et al., 2013). These algorithms involve two-step parameter estimation. First, the relative proportion of different subpopulations is estimated using the expression data of a set of cell type-specific reference genes, and then the estimated proportions are used to calculate the cell type-specific expression of other genes. These algorithms rely heavily on the fidelity of the genes chosen as cell type-specific reference genes. To circumvent this issue, Kang et al. (Kang, et al., 2019) developed a deconvolution method for the simultaneous estimation of cell-type proportions and cell type-specific gene expression from RNA-Seq data without using any reference gene information.

All these algorithms have been customized to analyze data of high-throughput experiments, like microarray and RNA-Seq. Though these high-throughput experiments are now accessible to most researchers, small to medium scale gene expression studies are still predominant where expressions of a handful of genes are measured by qPCR. Apart from having low experimental noise and ease of data analysis, qPCR for a handful of genes is easier to scale up for a large number of samples.

Gene expression is a dynamic process, and it changes with time and experimental conditions. Existing deconvolution methods do not consider such dynamics in gene expression while deconvoluting the data. Therefore, these algorithms are not suitable for the analysis of time-dependent experiments.

Keeping these limitations in mind, we have developed a computational tool DEBay that estimates cell type-specific gene expression profiles from qPCR data, given the proportions of different cell types in the population. DEBay can deconvolute both time-dependent and - independent gene expression data.

In contrast to existing regression-based deconvolution algorithms (Lahdesmaki, et al., 2005; Shen-Orr, et al., 2010), DEBay uses a Bayesian approach for model selection and parameter estimation. DEBay not only reports the expected relative gene expression in different cell types but also reports the probability distribution of these estimated parameters. This helps in understanding the uncertainty and quality of model selection and parameter estimation.

We have created an easy-to-use GUI for DEBay. It is implemented in Python and supported on MS Windows. DEBay would be useful in experiments where physical separation of pure cell-type is cumbersome, but proportions of different cell types in a sample are estimated through experimental techniques like quantitative image analysis, flow cytometry, Coulter counter.

## 2. Methods

### 2.1. Deconvolution algorithm

Let us consider a population of cells having multiple cell types or subpopulations. Experiments are performed using this heterogeneous cell population, and the expression of the target gene is measured in different samples by qPCR. Here, we consider two cases.

Case-1: The expression of the target gene in each cell type remains constant, but the population distribution of these cells varies among samples.

Case-2: We have time-dependent samples. Both the population distribution of cell types and the expression of the target gene in these cell types change with time.

Below, we discuss the mathematical formulations for both cases.

#### 2.1.1. Case-1

Consider a population of cells with *n* different cell types. Let’s assume that we have *m*+1 samples with varying proportions of these cell types. Let on an average each cell of the *k*^*th*^ type expresses *x*_*k*_ number of the target mRNA. The total number of the target mRNA in the population in sample *i* is,

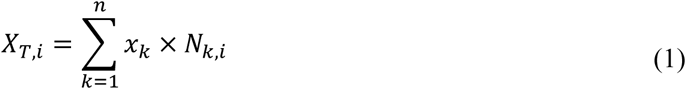

here, *i =* 0,1, 2, … *m* represent different samples, and the number of cells of *k*^th^ type in the *i*^th^ sample is *N*_*k,i*_. Total number of cells in the *i*^th^ sample, 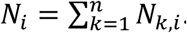. The sample *i* = 0 is the control samples (e.g., untreated cells) to be used for the fold-change estimation. Note that *x*_*k*_ is constant across all samples, but the population size of each cell type, *N*_*k,i*,_ changes.

The ΔΔ *C*_*t*_-method is used to measure normalized fold change in expressions of the target gene from quantitative PCR data (Kubista, et al., 2006; Livak and Schmittgen, 2001). Initially, fold change is calculated with respect to the control sample, followed by normalization with the reference/housekeeping gene. Here, we derive the relation between normalized fold change in expression of the target gene in the population and the expression of the target gene in each cell type.

The fold change in expression of the target gene in sample *i* is,

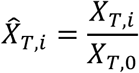

Here, *X*_*T*,0_ is the total number of mRNA of the target gene in the control sample.

Similarly, fold change in expression of the reference gene in sample *i* is,

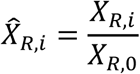

here, *X*_*R,i*_ is the total number of reference mRNA in sample *i* and *X*_*R*,0_ is the total number of reference mRNA in the control sample.

Normalized fold change in expression of the target gene in sample *i* is given by,

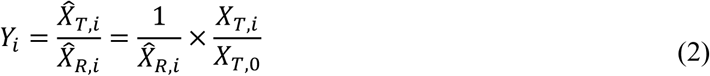

The ΔΔ *C*_*t*_ method estimates this fold change, *Y*_*i*_, from qPCR data.

From equation (1) and (2), we can write

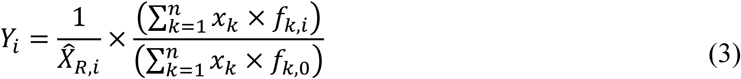

here, 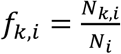 is the fractional population size of cell type *k* in sample *i*. Similarly, *f* _*k*,0_ is the fractional population size of *k*^th^ type cell in the control sample. The fractional population sizes of each cell type (*f*_*k,i*_) can be estimated from experiments like flow cytometry, quantitative image analysis.

Let’s define *ĝ*_*k*_, the Normalized Gene Expression Coefficient (NGEC) of cell type *k* as,

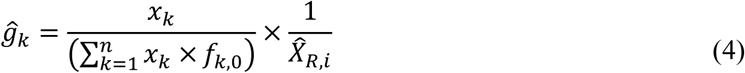

Now, Equation (3) can be written as,

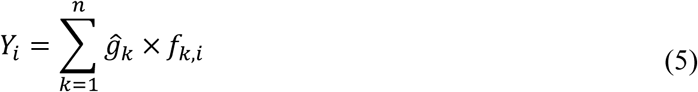

NGEC is a normalized measure of the level of expression of the target gene in the *k*^*th*^ cell type. In NGEC, the expression of the gene is normalized to the average expression of the gene across all cell types, and the fold change in expression of the reference gene.

Equation (5) should satisfy the constraint, *ĝ*_*k*_ ≥ 0. The values of *Y*_*i*_ and *f*_*k,i*_ for all *i*, are obtained from experiments. *ĝ*_*k*_ is the unknown parameter to be estimated from the data.

#### 2.1.2. Case 2

Here, we consider that we have samples collected for different time points (*i* = 0, *t*_1_, *t*_2_, …, *t*_*m*_), and the expression of the target gene in each cell type changes with time.

For a time-dependent system Equation (5) becomes,

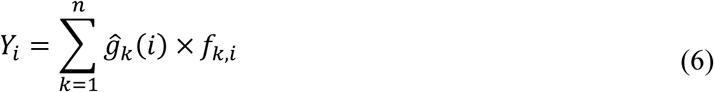

Here, *ĝ*_*k*_(*i*) is the time-dependent NGEC of the target gene in *k*^th^ type cell.

Commonly, the expression of a gene either increases or decreases or remains constant with time. Though complicated gene expression patterns, like oscillation, may be observed in some instances, we have considered that the time-dependent gene expression pattern in each cell-type follows one of the three predefined linear functions. We have used three predefined functions to reduce the complexity of the problem. These three functions are:

1. Linear time-dependent increase in gene expression:

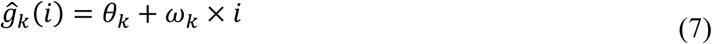 Here, *θ*_*k*_ and *ω*_*k*_ are constants and *θ*_*k*_, *ω*_*k*_ ≥ 0.
2. Linear time-dependent decrease in gene expression:

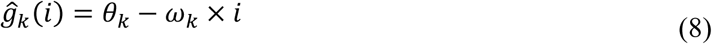

here *θ*_*k*_ and *ω*_*k*_ are constants, *θ*_*k*_, *ω*_*k*_ ≥ 0 *and θ*_*k*_ ≥ *ω*_*k*_ × *t*_*m*_.
3. Constant gene expression:

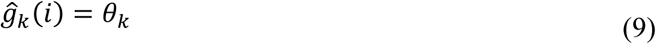

here *θ*_*k*_ is a constant and *θ*_*k*_ ≥ 0.

Here, we have considered time-varying gene expression. But the same mathematical formulation (Equation 6 to 9) can be used where gene expression in each cell type changes with the experimental condition, say a dose of a drug. In that case, *i* represents the dose of the drug, and *i* = 0 represent untreated cells. The mathematical formulation for NGEC and the algorithm used for its estimation remains the same as for the time-dependent problem.

### 2.2. Estimation of NGECs through Bayesian approach

We estimate the unknown NGEC of each cell-type through the Bayesian approach. Here, we consider the unknown parameters as a random variable and estimate the posterior probability distribution of the parameters given the experimental data. Using Bayes theorem (Kruschke, 2014) for our problem, we get

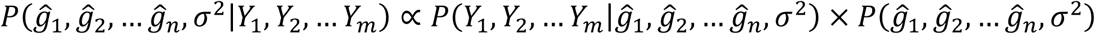

The term on the left-hand side is the posterior distribution of unknown parameters of our model. The first term on the right-hand side is the data likelihood, and the second term is the prior distribution of the parameters. *ĝ*_1_, *ĝ*_2,_ … *ĝ*_*n*_ are the NGECs of each cell-type that we want to estimate. *Y*_1_, *Y*_2,_ … *Y*_*m*_ are the normalized fold change in the expression of the target gene at various experimental conditions, estimated by qPCR. *σ*^2^ is the variance of the fold change in expression of the target gene. We assume that the variance is constant across all experimental conditions.

Using vector notation, the above equation can be written as,

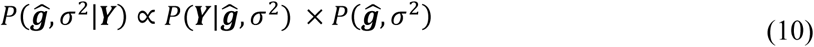

here, ***ĝ*** *=* (*ĝ*_1_, *ĝ*_2,_ … *ĝ*_*n*_); ***Y*** *=* (*Y*_1_, *Y*_2,_ … *Y*_*m*_). *P*(***ĝ***, *σ*^2^|***Y***) is the posterior distribution of the unknown parameters. *P*(***Y***|***ĝ***, *σ*^2^) is the data likelihood. *P*(***ĝ***, *σ*^2^) is the prior distribution of the unknown parameters.

In this formulation, both ***ĝ*** and *σ*^2^ are unknown parameters. We assume that the experimental observations (***Y***) are Normally distributed around the true mean with some unknown variance (*σ*^2^) (Kruschke, 2014; Raue, et al., 2011; Raue, et al., 2013). For *m*+1 different samples and *n* different cell types, the data likelihood is,

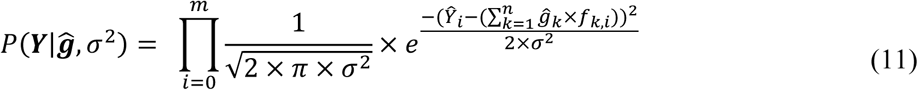

The NGECs cannot be negative. Therefore, for parameter estimation, we have used a truncated normal prior distribution (0 to +∞) for ***ĝ*** and an inverse gamma prior distribution for the variance (Gelman, et al., 2013). We have employed a hierarchical Bayesian approach where the prior distributions of the NGECs are sampled from a hyperprior (Kruschke, 2014). We made use of conjugate-prior to the Normal data likelihood, as described by Murphy (Murphy, 2007) and Clyde et al. (Clyde, et al.). The priors are defined as,

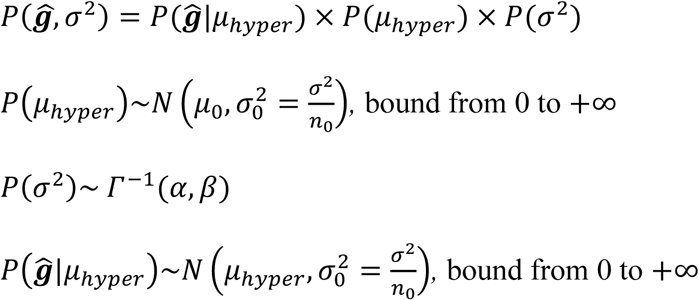

*P*(*μ*_*hyper*_) is the distribution of the hyperprior from which the prior distributions of ***ĝ*** are sampled. *α* and *β* are the parameters that control the height and width of the inverse gamma distribution, respectively. *n*_0_ is the scale parameter that controls the variance in the prior 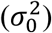 relative to the variance in the data (*σ*^2^). In the GUI of DEBay, the user can define *α, β, μ*_0,_ and *n*_0_.

The posterior distribution is estimated by Markov Chain Monte Carlo (MCMC) using the NUTS sampler (Betancourt, 2017; Hoffman and Gelman, 2014). Complete estimation steps were implemented in Python through the PyMC3 package (Salvatier, et al., 2016).

For the time-dependent system discussed in section 2.1.2, we have defined the gene expression coefficient as a function of time, in terms of unknown parameters (***θ, ω***). The priors are defined as,

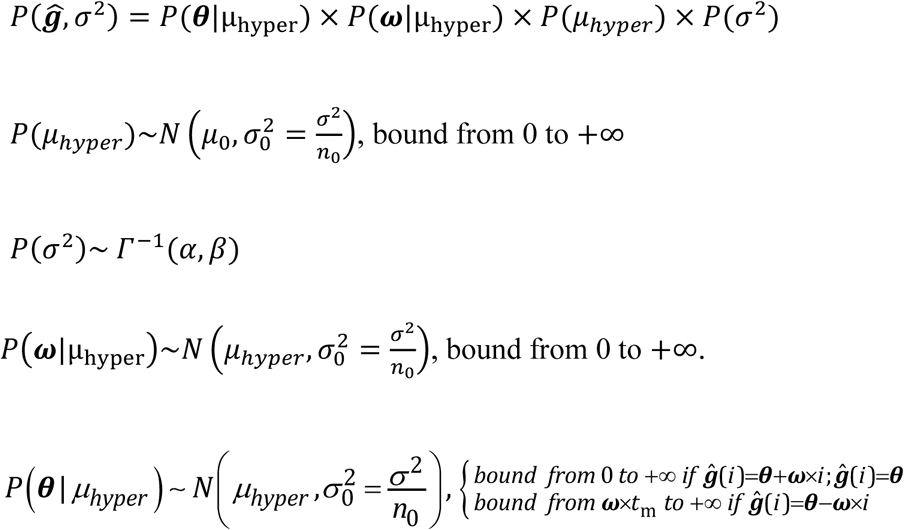

For the time-dependent system, the data likelihood is evaluated at the observed time points. For *m* discrete time points and *n* different cell types, the data likelihood is,

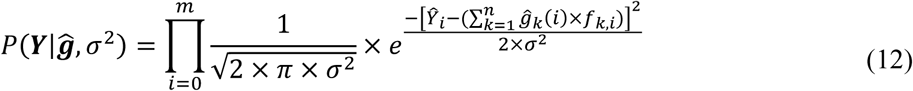

Here, *ĝ*_*k*_(*i*) is evaluated as a function of time (*i*) at the observed time points. *ĝ*_*k*_(*i*) can follow any of the three predefined functions (Equation (7)-(9)). In general, for *n* cell types and three different predefined functions, we have 3^*n*^ possible combinations. Each function combination is a possible model, and we estimate the posterior distribution for all models given the experimental data. The optimal model is selected based on the Bayes Information Criterion (BIC) (Lorah and Womack, 2019; Vrieze, 2012),

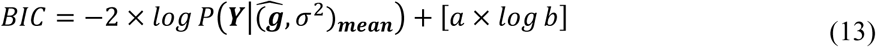

here 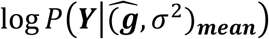 is the log-likelihood evaluated at the mean of each parameter distribution; *a* is the number of unknown parameters, including the variance and the hyperprior; *b* is the number of observed data points. The model with minimum BIC is considered as the optimal model.

From the optimal model, we get the distribution of each parameter in the predefined functions (Equation (7)-(9)). By using random numbers from these parameter distributions in *ĝ*_*k*_(*i*), we get the distribution of the NGEC of each cell-type for each discrete time point.

### 2.4. Real-Time PCR

We have evaluated DEBay with both synthetic and experimental data. Experimental data was generated by real-time PCR. Total RNA was isolated using TRI reagent (Sigma, St. Louis, MO, USA). Genomic DNA contamination was removed by DNAse treatment. cDNAs were prepared from an equal amount of total RNA from all samples using Verso cDNA synthesis kit (Thermo Fisher Scientific, Waltham, MA, USA). Real-time PCR was performed using PowerUp SYBR Green (Thermo Fisher Scientific, Waltham, MA, USA) in Rotor-Gene Q real-time PCR cycler (QIAGEN, Hilden, Germany). Cyclophilin A was used as the reference gene, and the experiments were performed in triplicates. Data analysis was done using LinRegPCR (Ramakers, et al., 2003).

## 3. Results

### 3.1. Deconvolution of qPCR data to estimate cell type-specific gene expression

DEBay takes two data - a) fold change in expression of a target gene in samples, and b) proportion of each cell-type in these samples. Using these data, DEBay estimates the relative expression of the gene in each cell type. The expression of a gene can be time-dependent or independent. DEBay handles both cases.

DEBay estimates the relative expression of a gene in a particular cell type as Normalized Gene Expression Coefficient (NGEC), using the Bayesian method of parameter estimation. The definition of NGEC, it’s mathematical formulations, details of the algorithm, and the estimation strategy are given in the Methods section. Figure 1a outlines the workflow of the estimation strategy.

**Figure 1:**
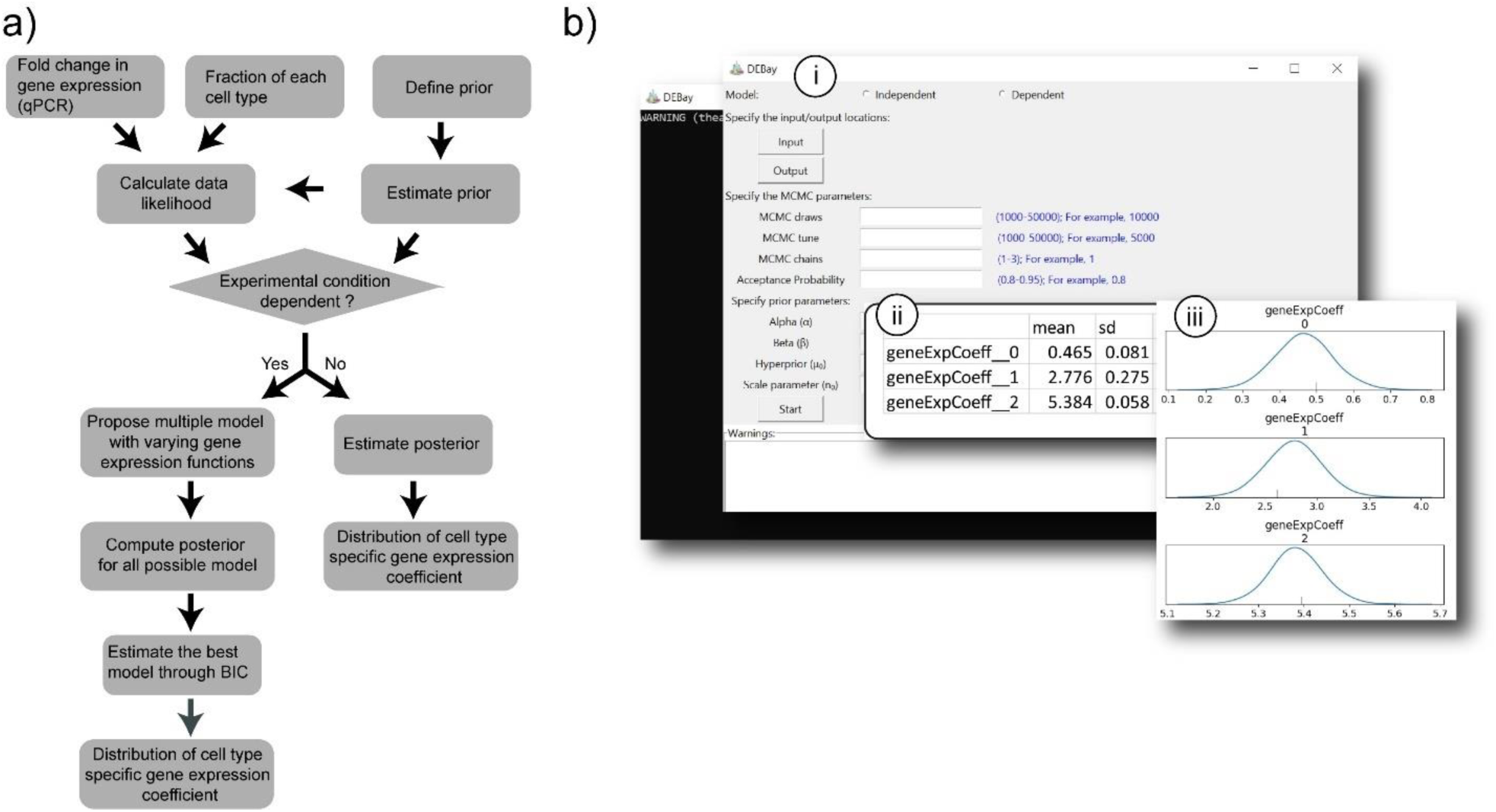
a) Workflow of the algorithm. b) Snapshots of the GUI (*i*) and outputs (*ii* and *iii*) generated by DEBay.

DEBay takes the following inputs from the user- a) whether to use time-dependent or - independent model, b) data in an MS Excel file, c) parameter values for MCMC, d) parameter values specifying the prior distributions. The estimated average values of NGECs are saved as a tab-delimited text file along with the statistics. The probability distributions of the NGECs are saved in a graphical format.

DEBay is implemented in Python. We have created a GUI for DEBay. Glimpses of the GUI and data output are shown in Figure 1b. A standalone windows installer for DEBay and a user manual is available at SourceForge (https://sourceforge.net/projects/debay). The python packages used in DEBay uses C libraries for faster computation. Therefore, a GCC would help to improve the performance of DEBay but it is not mandatory. MinGW, a complete run time environment for GCC for Windows is available at SourceForge (https://sourceforge.net/projects/mingw-w64/).

### 3.2. Evaluation of the algorithm

We have used both synthetic data and experimental data to evaluate DEBay. Detailed information about the generation of the synthetic data is given in the Supplementary Text.

#### 3.2.1. Case 1

The assumption here is that the expression of the target gene in a cell type does not vary among the sample, but the population size of each cell-type varies. We generated 1000 different sets of synthetic data for a population of cells having four types of cells. These data sets were generated by varying the population size and the mean number of mRNAs for each cell type. The detailed information about the synthetic data generation is given in the Supplementary Text.

Figure 2 shows the analysis of a representative synthetic data set using DEBay. There were five samples (S1 to S5) with different population distribution for four cell-types (Figure 2a). Figure 2b shows the gene expression data for each sample. Data of Figures 2a and 2b were used as inputs for DEBay to estimate NGECs for each cell type. As shown in Figure 2c, the estimated mean NGECs for all the cell types (Figure 2c) are close to the respective actual values (Figure 2d) calculated algebraically using Equation (4). Figure 2e shows the distribution of the estimated cell-type-specific NGECs.

**Figure 2:**
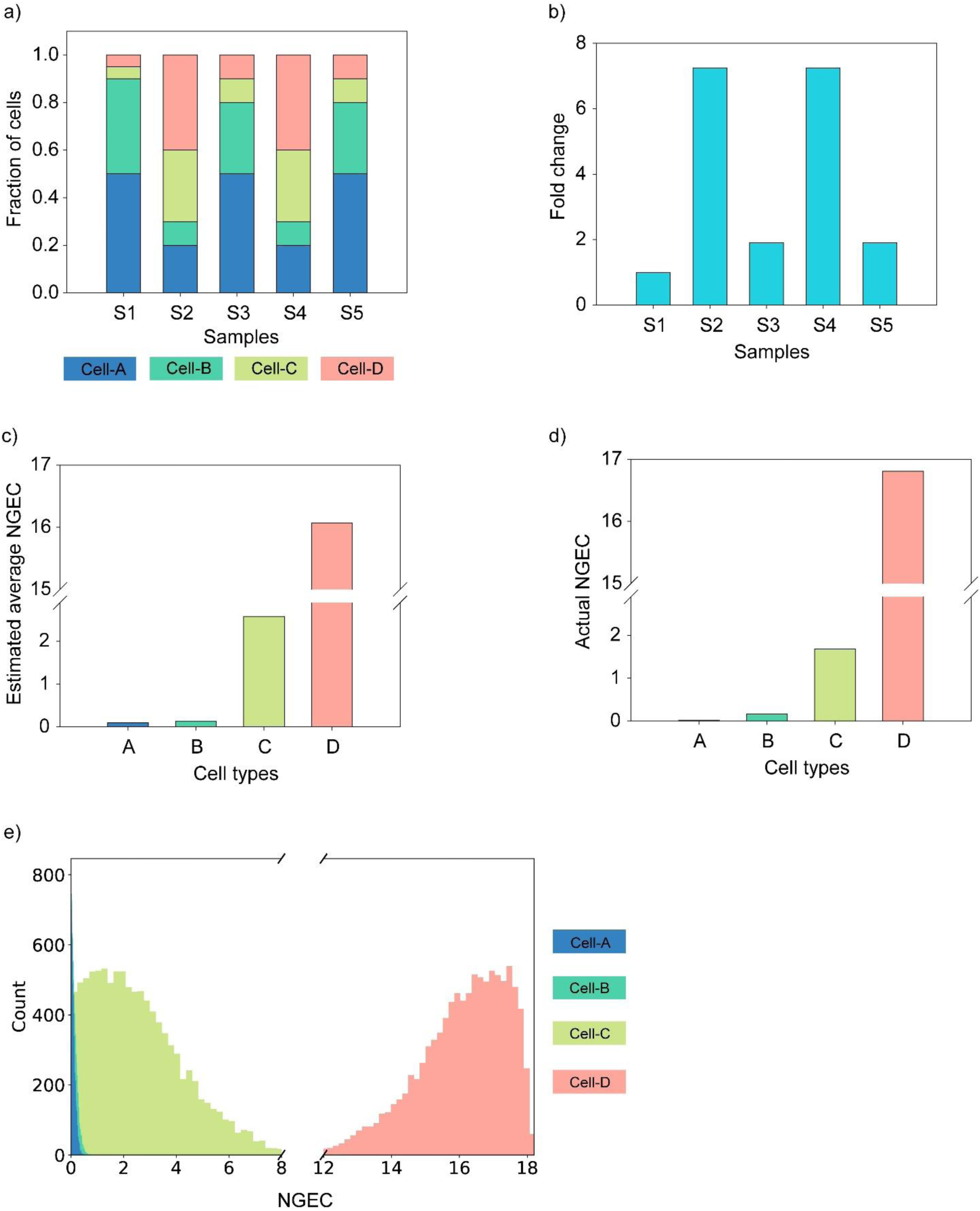
Deconvolution of synthetic data using DEBay. This data set has five samples, and each sample is composed of four types of cells. The population size of each cell type varies among samples (a). (b) shows the fold change in expression of the target gene in different samples. The estimated average NGECs for different cell types are shown in (c). The actual values of the NGECs are shown in (d). e) Distribution of the estimated NGECs of four cell types.

Supplementary Figure S2a shows the deviation of the estimated NGECs from the actual ones, relative to the standard deviation of the estimated NGECs, for all the thousand synthetic data sets. In all cell-types, the distribution of the deviation showed a tight cluster around zero. Most of the actual NGECs were within one standard deviation of the estimated NGECs (Supplementary Figure S2a). The actual and estimated NGECs showed a strong positive correlation with r^2^ = 0.99 (Supplementary Figure 2b).

We generated real data by a qPCR experiment to evaluate DEBay. We used three different human breast cancer cell lines, MCF-7, MDA-MB-231, and MDA-MB-468. These cell line expresses different amounts of EGFR1 (MDA-MB-468 > MDA-MB-231 > MCF-7) (Davidson, et al., 1987). We have measured the expression of EGFR1 in all three cells. The *Ct* values of EGFR1 expression were normalized to the reference gene (Cyclophilin A). The normalized *Ct* values of MDA-MB-468, MDA-MB-231, and MCF-7 were 0.9, 1.1, and 1.4, respectively. Higher the *Ct* value, lower is the expression level.

We mixed these three cell lines in various proportions to prepare three samples S1, S2, and S3 (Figure 3a). Through qPCR, we measured the fold change in EGFR1 expression in samples S2 and S3 with respect to S1 (Figure 3b). Subsequently, this fold change data were deconvoluted using DEBay.

**Figure 3:**
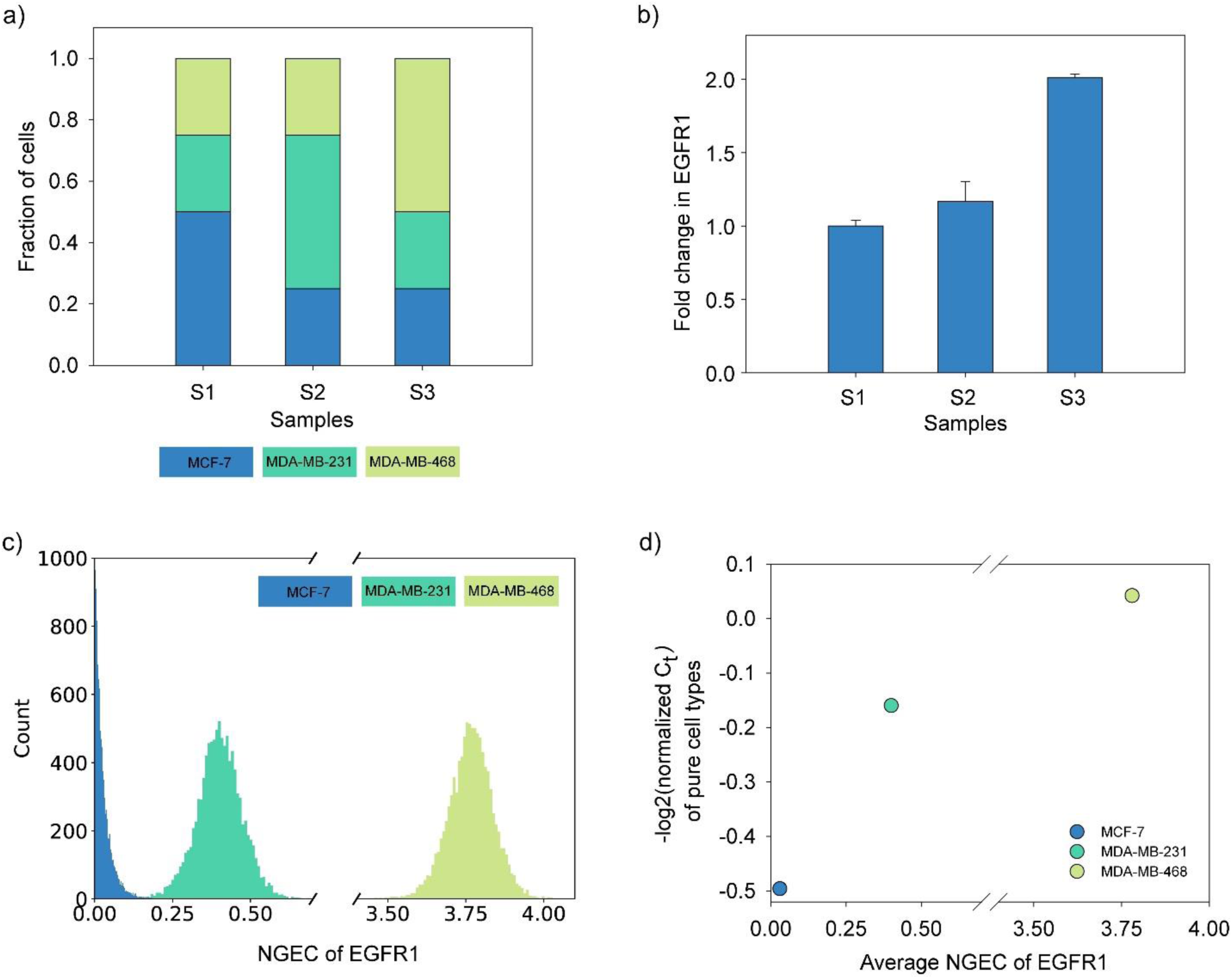
Deconvolution of real qPCR data using DEBay. (a) Three samples were prepared by mixing different proportions of MCF-7, MDA-MB-468, and MDA-MB-231 cells. (b) shows normalized fold change in expression of EGFR1 in different samples as measured by qPCR. c) Distribution of estimated NGECs of EGFR1 expression in three cell types. d) shows the correspondence between the experimentally measured level of expression of EGFR1 in three pure cell lines and NGECs of these cells estimated by DEBay from three mixed samples. *Ct* values of EGFR1 were normalized with the *Ct* values of cyclophilin A. Normalized *C*t value is a proxy of gene expression level and is inversely related to the level of expression of a gene. Negative log2 transformation has been used for better visualization of the data.

Figure 3c shows the estimated distribution of NGECs for EGFR1 in three pure cell-lines. NGECs for EGFR1 expression in these cell lines have the same pattern, MDA-MB-468 > MDA-MB-231 > MCF-7, as observed by qPCR. Figure 3d shows the correspondence between experimentally determined *Ct* values and the estimated NGECs.

#### 3.2.2. Case 2

Here we have considered that both the population distribution and the expression of the target gene in each cell-type change with time. We have used three sets of synthetic data to evaluate the algorithm. The detailed information about the synthetic data generation is given in the Supplementary Text.

As a representative data, the results of synthetic data of set-1 are shown in Figure 4, and the rest are available in supplementary text (Supplementary Figure S3 and S4). Here, we have samples for five-time points, *t* = 0 to 48. Each sample is a mixture of four cell types. Figures 4a shows the changing proportions of these cell types with time. The fold change in expression of the target gene with time is shown in Figure 4b. This fold change data were deconvoluted using DEBay.

**Figure 4:**
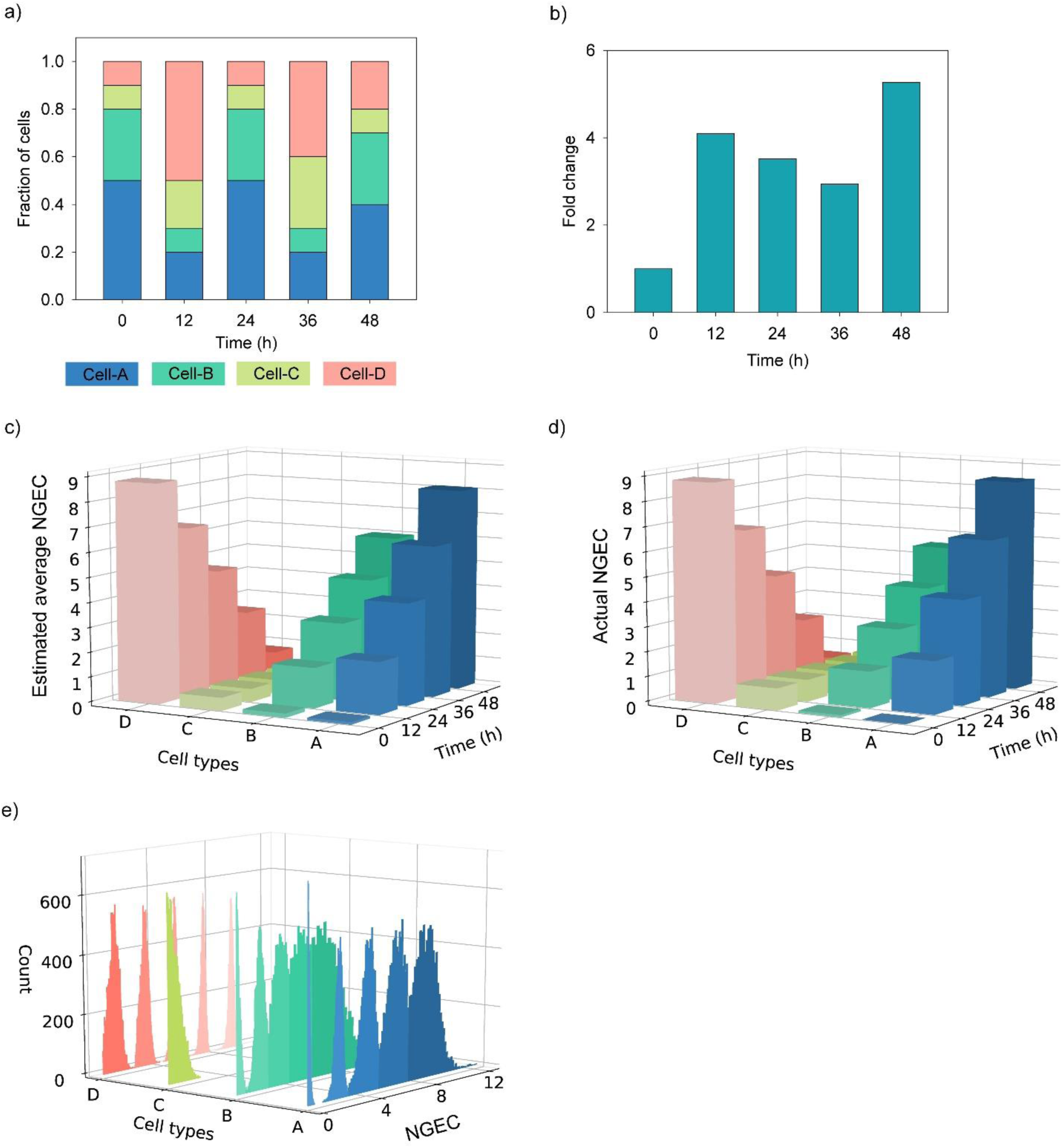
Deconvolution of synthetic data for time-dependent gene expression using DEBay. This data set has five samples corresponding to five time-points. Each sample is composed of different proportions of four types of cells. (a) shows the change in the proportions of cell types with time. (b) Fold change in expression of the target gene in the whole population with time. Deconvolution was performed using the time-dependent model of DEBay. The estimated average NGECs and the actual NGECs for different cell types are shown in (c) and (d), respectively. e) Distribution of the estimated time-dependent NGECs. Time points are represented by increasing order of color intensities. The lowest intensity denotes the initial time point, and the highest intensity denotes the end time point.

For the time-dependent case, DEBay considers three linear models for each cell type (discussed in Section 2.1.2) and reports the best combination of models. Figure 4c shows the estimated average time-dependent NGECs for four cell types for this data set. As per this deconvolution, with time, expression of the gene increases in cell type A and B and decreases in cell type D (Figure 4c). Expression of the gene remains constant in cell type C. Similar behavior is observed when NGECs were calculated algebraically from the synthetic data (Figure 4d). Figure 4e shows the distribution of the estimated NGECs as a function of time.

Supplementary Figure S5a shows the deviation between the actual and the estimated NGECs for all three synthetic data sets. In most of the cases, the actual NGECs lies within 1.5 standard deviations of the estimated NGECs. The estimated NGECs showed a strong positive correlation to the actual NGECs with r^2^ = 0.97 (Supplementary Figure S5b).

Subsequently, we analyzed an experimental data set using DEBay. We had earlier investigated the dynamics of EGF-induced Epithelial to Mesenchymal Transition (EMT) of MDA-MB-468 cells (Devaraj and Bose, 2019). We had observed that a population of MDA-MB-468 cells had three subpopulations having different morphologies-cobble, spindle, and circular. It was observed that the cobble cells were non-migratory and epithelial-like, whereas spindle and circular cells were migratory and mesenchymal-like (Devaraj and Bose, 2019). Treatment with EGF induced EMT in MDA-MB-468 cells, and the proportions of three cell types changed, both with time and dose of EGF (Devaraj and Bose, 2019).

For a time-dependent experiment, we measured the population distribution of three cell-types in EGF-treated MDA-MB-468 cells at different time points using quantitative image analysis (Devaraj and Bose, 2019). Fold change in expression of Vimentin and Snail1, two markers of EMT, was estimated by qPCR. Figures 5a and b show the data from this experiment.

**Figure 5:**
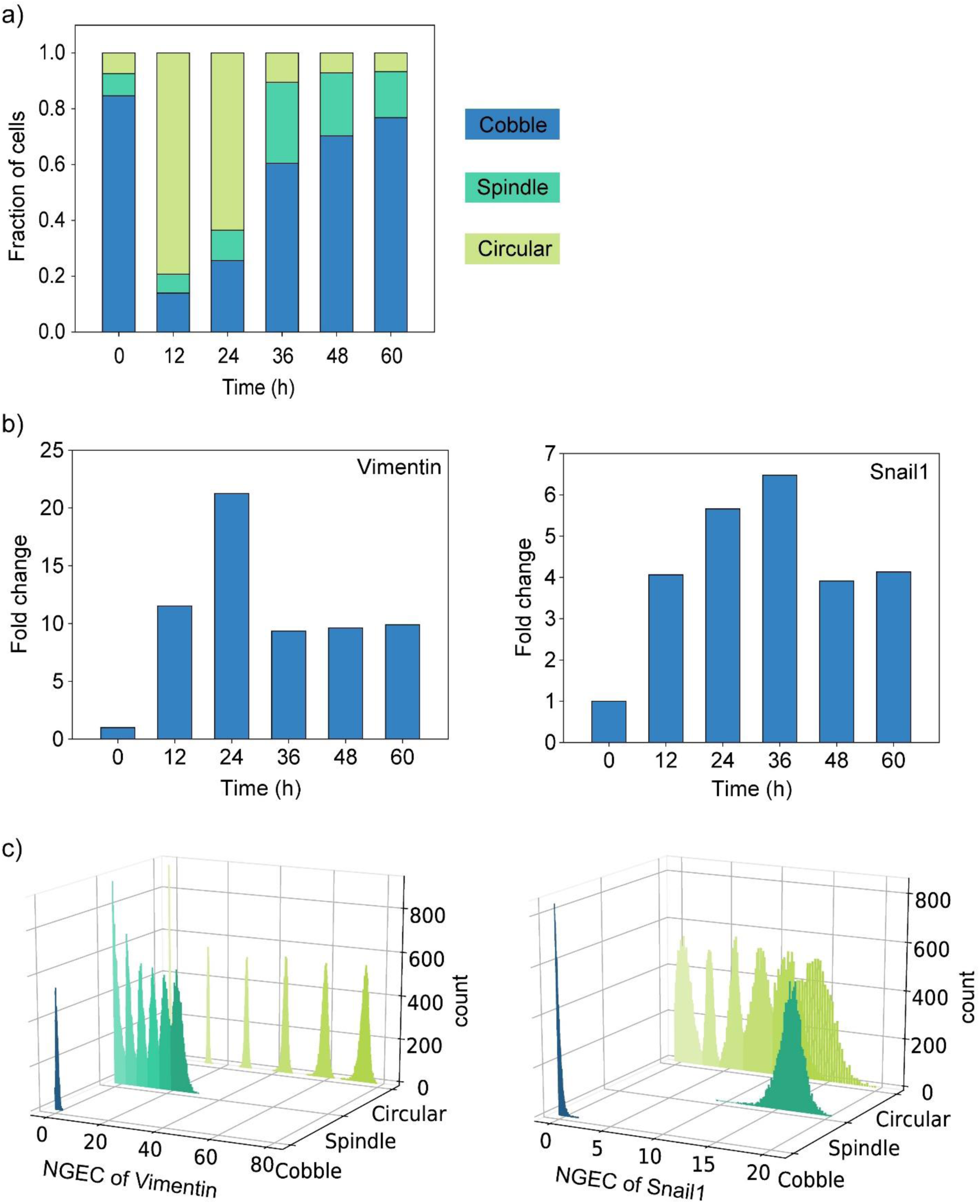
Deconvolution of a time-dependent gene expression data using DEBay. MDA-MB-468 cells were treated with EGF for different durations, and the proportions of three cell types were measured by quantitative image analysis (a). qPCR was used to estimate the time-dependent changes in the expression of two markers of EMT, Vimentin, and Snail1(b). These data were deconvoluted using the time-dependent model of DEBay. c) Time-dependent distribution of NGEC of Vimentin and Snail1, in three cell types. Time points are represented by increasing order of color intensities. The lowest intensity denotes the initial time point, and the highest intensity denotes the end time point.

The qPCR data were deconvoluted using DEBay to estimate the time-dependent changes in NGECs of Vimentin and Snail1 for three cell types (Figure 5c). The NGECs of Vimentin and Snail1 for cobble cells are extremely low and do not change with time. On the other hand, NGECs of these two genes for the spindle and circular cells are either very high or increased with time (Figure 5c). It is known that the expression of Vimentin and Snail1 is low in epithelial cells and high in mesenchymal cells (Lamouille, et al., 2014; Zeisberg and Neilson, 2009). Therefore, from the NGEC values, we can say that the spindle and circular cells are possibly mesenchymal cells, and cobble cells are epithelial. This observation matches the migration behavior of these cell types.

## 4. Discussion

In this work, we have presented DEBay, a tool for computational deconvolution of quantitative PCR data to estimate time-dependent and -independent expression of a target gene in different subpopulations in an ensemble of cells. The time-dependent model of DEBay can also be used to deconvolute data where gene expression in subpopulations changes with the experimental condition like the dose of a drug.

DEBay estimates the relative level of expression of a gene in a subpopulation as the Normalized Gene Expression Coefficient (NGEC). NGEC is mathematically derived from the fold change in gene expression measured by qPCR. In essence, NGEC is a proxy of the normalized gene expression level in each cell type in the sample.

The existing deconvolution algorithms consider that the gene expression of a cell type in a population remains constant with time or across experimental conditions. DEBay addresses this problem by considering three different models of time-dependent gene expression – linear increase, linear decrease, and constant. One could envisage various types of nonlinear gene expression patterns, and the algorithm of DEBay can be easily altered to accommodate such dynamics. However, an increase in the number of alternative models increases the complexity of the problem and computation time. Further, nonlinear models suffer from the problem of overfitting when the number of data points is low. Therefore, for the current version of DEBay, we have used three linear models that are most commonly observed in experiments.

Most of the deconvolution algorithms uses the frequentist approach to estimate the unknown parameter. These methods converge to a point estimate of the unknown parameter and usually report *P* value or confidence interval of the estimated parameter. This approach do not address the probability distribution of the estimated parameters. In our method, we have used the Bayesian method of parameter estimation. In this approach, the parameters are considered as random variables, and we estimate the posterior distribution of the parameters based on the observed data. Through this approach, we can estimate the credible interval of the estimated parameters.

In our method, we have used a hierarchical model structure. The prior distributions of cell-type-specific gene expression coefficients are defined based on hyperprior. Through this model structure, each cell-type-specific coefficient is indirectly constrained by all the observed data through hyperprior. The credible intervals of cell-type-specific coefficients are pulled towards the mode of the hyperprior. Thereby, increasing the efficiency of the sampling (Kruschke, 2014).

We have evaluated DEBay with real biological data and synthetic data sets. Our algorithm performed reasonably well in all cases. DEBay has been developed, keeping in mind the needs of low-throughput but widely used qPCR experiments. The GUI of DEBay is intuitive and straightforward. One can use the fold change data obtained from qPCR experiments without any correction or transformation. The output files created by DEBay are also self-explanatory.

## Acknowledgments

We are thankful to the DBT Programme Support Facility at IIT Guwahati, funded by the Department of Biotechnology, Government of India (Project No. BT/PR13560/COE/34/44/2015) for resources and research facilities.

## Notes

### Competing Interest Statement

The authors have declared no competing interest.

