## Supplementary information for "DEBay: a computational tool for deconvolution of quantitative PCR data for estimation of cell type-specifc gene expression in a mixed population"

### S1. Synthetic data to evaluate the algorithm

We have used experimental data as well as synthetic data to evaluate the deconvolution algorithm. To generate synthetic data, we considered five samples of ($i$ = S1, S2, S3, S4, and S5) having $N = 10^6$ cells per sample, with four subpopulations, A, B, C, and D.

In general, suppose for the $i^{th}$ sample, we have $N_{k,i}$ number of $k$-type cells. $N_{k,i} = N \times f_{k,i}$, where $f_{k,i}$ is the fractional size of the $k^{th}$ subpopulation in the $i^{th}$ sample. The total number of mRNAs of the target gene in $k^{th}$ subpopulation in this sample, $X_{k,i} = \sum_{j=1}^{N_{k,i}} x_{j,k,i}$. Here, $x_{j,k,i}$ is the number of mRNAs of the target gene in the $j^{th}$ cell of $k$-type in $i^{th}$ sample. Therefore, the total number of the target mRNAs in the whole population is $\sum_k X_{k,i}$.

The fold change in target gene expression in the $i^{th}$ sample was calculated with respect to the control sample. We have considered $i$ = S1 as the control sample. Therefore, the fold change in expression of the target gene in the $i^{th}$ sample is

$$Y_i = \frac{\sum_k X_{k,i}}{\sum_k X_{k,S1}} = \frac{\sum_k \sum_{j=1}^{N_{k,i}} x_{j,k,i}}{\sum_k \sum_{j=1}^{N_{k,S1}} x_{j,k,S1}} \qquad (1)$$

Usually, the change in the expression of the target gene is reported as normalized fold change, where the normalization is done with respect to the fold change in the expression of a reference/housekeeping gene. For simplification, we have considered that normalization term as one. Therefore, Equation (1) in itself gives us the normalized fold change in the target gene in the $i^{th}$ experimental condition.

We varied the fractional population size of each cell-type ($f_{k,i}$) such that $\sum_k f_{k,i} = 1$. The number of target mRNAs in each $k^{th}$ type cell ($x_{j,k,i}$), in a sample, was generated by repeated sampling

($N_{k,i}$ times) from a Normal distribution $N(\mu_{k,i}, \sigma_{k,i})$. The number of copies of mRNAs of highly expressed genes in human cells varies in the range of $10^3$-$10^5$ per cell (*1*). Therefore, $\mu_{k,i}$ was varied from $10^2$-$10^5$.

The noise in gene expression is often quantified as $\eta_{k,i} = \frac{\sigma_{k,i}}{\mu_{k,i}}$. If we fix the level of the noise ($\eta_{k,i}$), then $\sigma_{k,i}$ can be calculated for a given $\mu_{k,i}$. For simplicity, we considered equal noise in all the subpopulations in a sample. We generated samples with different level of noises and calculated the fold change in expression using Equation (1). We observed that the noise in mRNA level did not have any effect on the calculated fold change (Figure S1). The effect of noise is nullified since we considered constant noise across all cell types in a population. Therefore, in all further data sets, we used $\sigma_{k,i} = 10^{-1}\times \mu_{k,i}$.

**S2. Evaluating DEBay with time-independent gene expression data**

We generated 1000 independent synthetic data sets. For a particular data set, the population size of each cell-type ($f_{k,i}$) varied randomly among the samples, but the mean level of expression of the target gene ($\mu_{k,i}$) remained the same ($\mu_{k,i}$ = constant for all $i$). We varied $f_{k,i}$ by sampling from a uniform distribution U(0,1) such that $\sum_k f_{k,i} = 1$. The mean number of target mRNAs in each cell type, $\mu_k$ was varied randomly from $10^2$-$10^5$. Accordingly, $\sigma_k$ was decided such that $\sigma_k = 10^{-1}\times \mu_k$. As mentioned earlier, repeated sampling was done from $N(\mu_{k,}, \sigma_{k,})$ to generate the data for each cell in each subpopulation. These sampled data were used to calculate the population-level fold change in expression using Equation 1. The population-level synthetic data were deconvoluted using DEBay to estimate the Normalized Gene Expression Coefficient (NGEC) of each cell type.

**S3. Evaluating DEBay with time-dependent gene expression data**

In time time-dependent case, we have used three sets of synthetic data sets. Each data set has five samples, representing five time points, from 0 to 48 h at an interval of 12 h. From equation 1, the fold change in the target gene expression is,

$$Y_i = \frac{\sum_k X_{k,i}}{\sum_k X_{k,0}} = \frac{\sum_k \sum_{j=1}^{N_{k,i}} x(i)_{k,j}}{\sum_k \sum_{j=1}^{N_{k,0}} x(0)_{k,j}} \tag{2}$$

here $i = 0, 12, 24, 36, 48\ h$. $x(i)_{k,j}$ is the time-dependent function representing the number of target mRNAs in the $j^{th}$ cell of type $k$ at the $i^{\text{th}}$ time point. We considered four cell types, $k$ = A, B, C, and D.

The functions used to generate the number of target mRNAs are given in the Supplementary Table S1a-c. The parameters in the functions ($p = (\theta, \omega)$) are generated using random numbers from a normal distribution, $N \sim (\mu_p, \sigma = 10^{-1} \times \mu_p)$. The values of $\mu_\theta, \mu_\omega$ are given in the Supplementary Table S1a-c. The fraction of each cell type is varied from 0 to 1, such that the summation of fractions of all cell types at a given experimental condition is 1. Using the fractions of different cell types and the mRNAs expression in each cell type in equation (2), the population-level fold change in target gene expression is calculated. This population-level fold change data were deconvoluted using DEBay.

**Table S1:**

**a) For data set-1**

| **Cell-types** | **mRNAs as a function of time** | $\mu_\theta$ | $\mu_\omega$ |
|---|---|---|---|
| A | $\theta + \omega \times i$ | 100 | 2000 |
| B | $\theta + \omega \times i$ | 1000 | 1300 |
| C | $\theta$ | 10000 | - |
| D | $\theta - \omega \times i$ | 100000 | 2000 |

**b) For data set-2**

| **Cell-types** | **mRNAs as a function of time** | $\mu_\theta$ | $\mu_\omega$ |
|---|---|---|---|
| A | $\theta - \omega \times i$ | 15000 | 250 |
| B | $\theta$ | 1000 | - |
| C | $\theta$ | 500 | - |
| D | $\theta + \omega \times i$ | 5000 | 100 |

**c) For data set-3**

| **Cell-types** | **mRNAs as a function of time** | $\mu_\theta$ | $\mu_\omega$ |
|---|---|---|---|
| A | $\theta$ | 2500 | - |
| B | $\theta + \omega \times i$ | 500 | 200 |
| C | $\theta - \omega \times i$ | 12500 | 300 |
| D | $\theta$ | 100 | - |

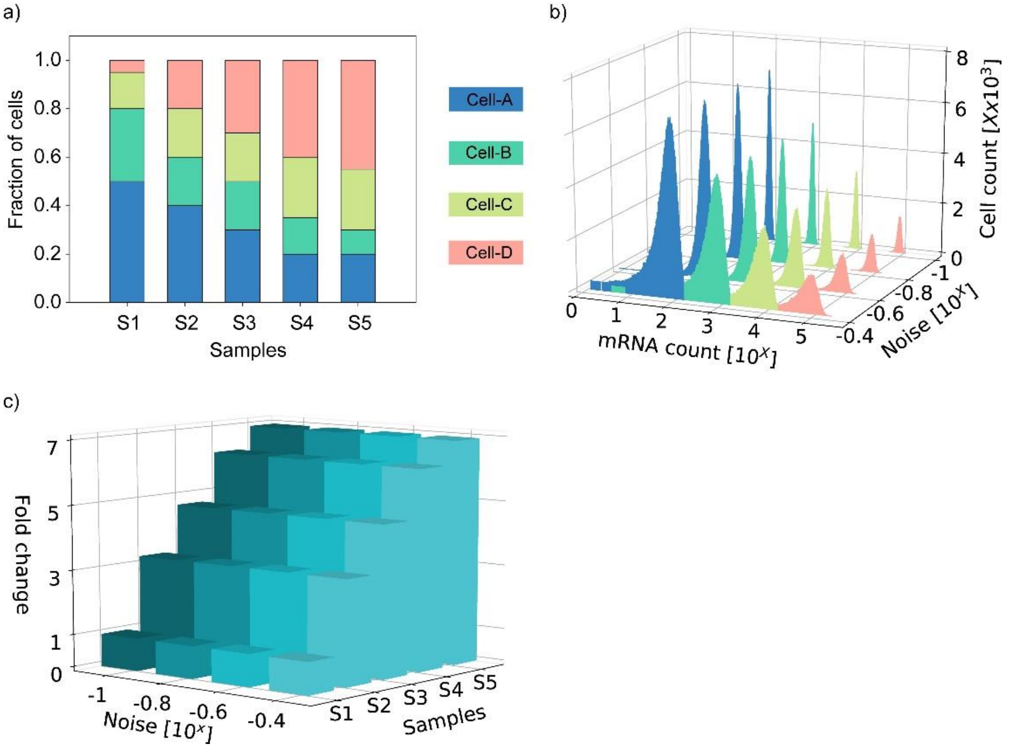


Figure S1: Effect of noise in mRNA level on the estimation of population-level fold change in expression. (a) Each sample was a mixture of four types of cells (A, B, C, and D). The mean number of mRNAs in each cell type was fixed. The number of mRNA in each cell was sampled from distributions having four different levels of noises, $\eta = \sigma/\mu$. (b) shows the sampling distribution for sample S1. (c) Fold change in gene expression was calculated from these randomly sampled mRNA data.

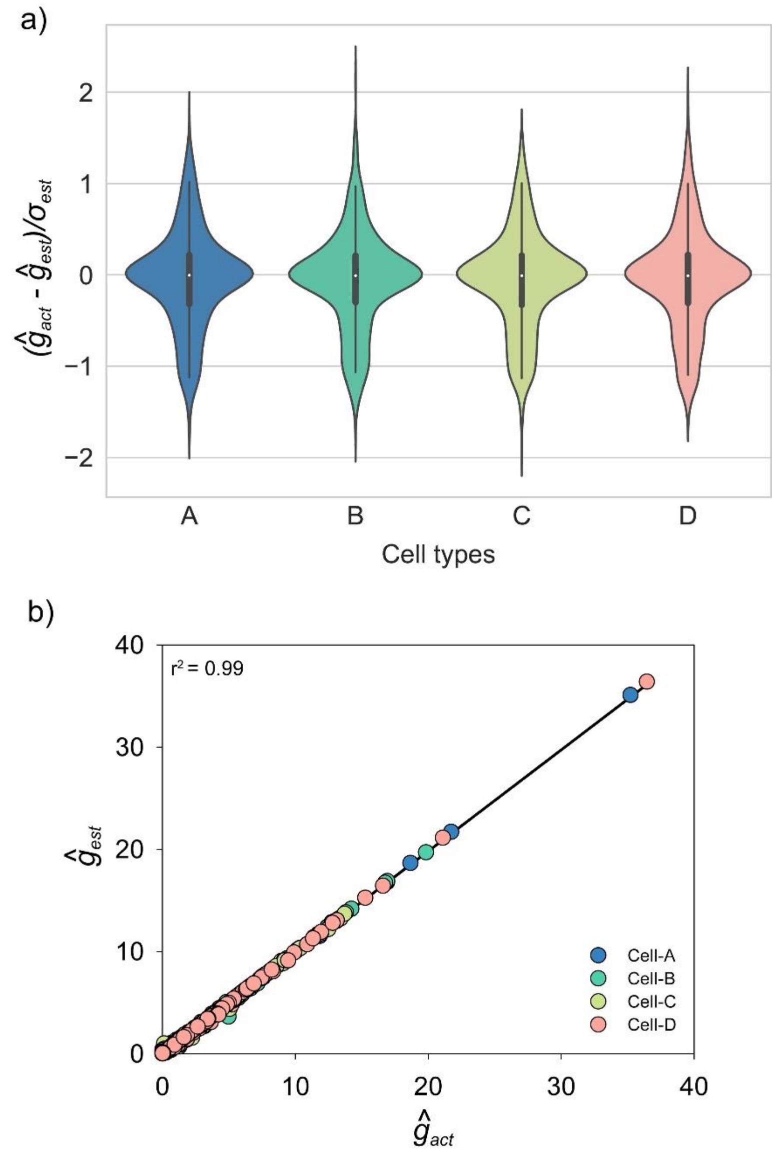


Figure S2: Evaluation of the performance of DEBay for synthetic data sets. One thousand independent synthetic data sets were generated with different population fractions and different mean levels of expression in each of the four cell-types. a) The deviation between the mean of estimated NGECs ($\hat{g}_{est}$) and the actual NGECs ($\hat{g}_{act}$) to the standard deviation of the estimated NGECs ($\sigma_{est}$). b) Correlation between the mean of estimated NGECs ($\hat{g}_{est}$) and the actual NGECs ($\hat{g}_{act}$).

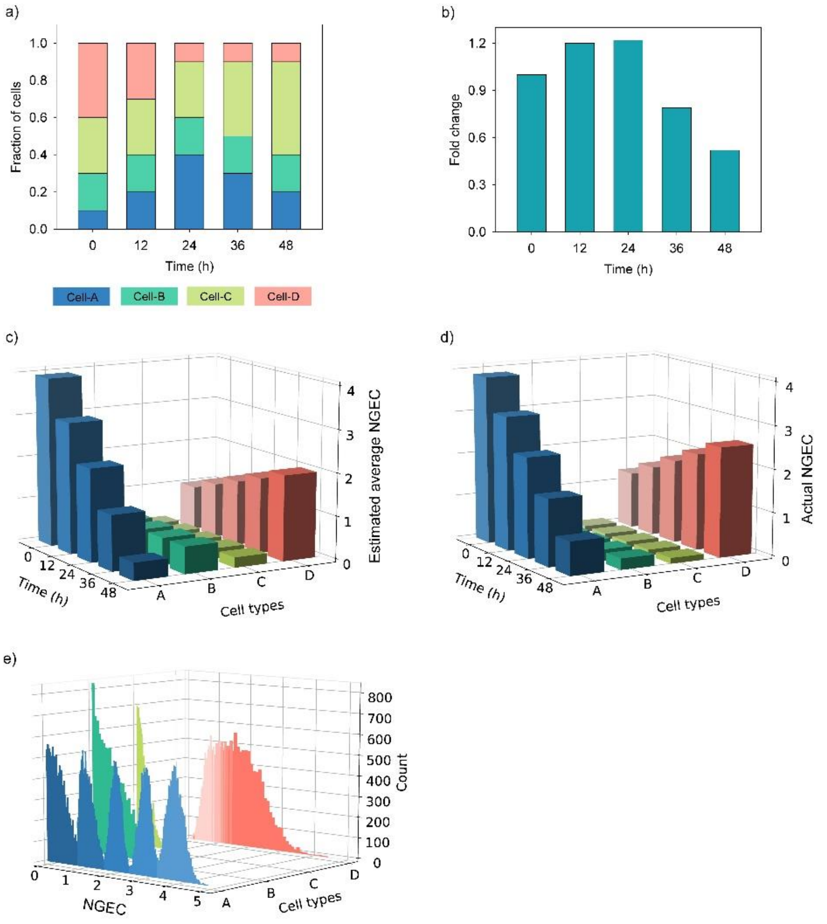


Figure S3**:** Deconvolution of synthetic data (Set -2) for time-dependent gene expression using DEBay. This data set has five samples corresponding to five time-points. Each sample is composed of different proportions of four types of cells. (a) shows the change in the proportions of cell types with time. (b) Fold change in expression of the target gene in the whole population with time. Deconvolution was performed using the time-dependent model of DEBay. The estimated average NGECs and the actual NGECs for different cell types are shown in (c) and (d), respectively. e) Distribution of the estimated time-dependent NGECs. Time points are represented by increasing order of color intensities. The lowest intensity denotes the initial time point, and the highest intensity denotes the end time point.

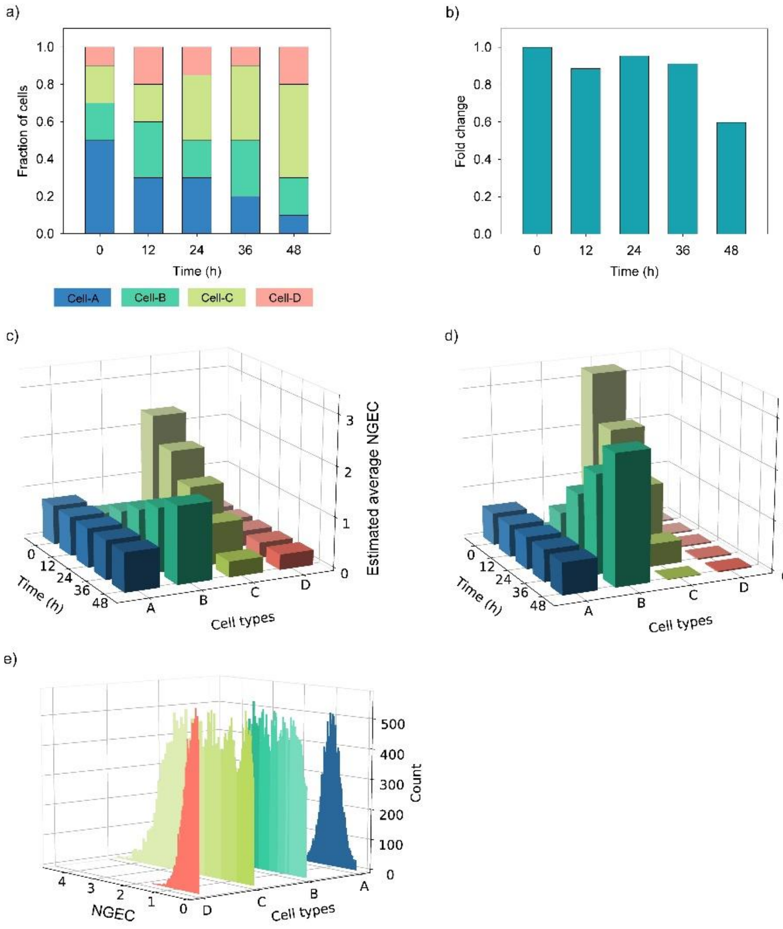


Figure S4: Deconvolution of synthetic data (Set -3) for time-dependent gene expression using DEBay. This data set has five samples corresponding to five time-points. Each sample is composed of different proportions of four types of cells. (a) shows the change in the proportions of cell types with time. (b) Fold change in expression of the target gene in the whole population with time. Deconvolution was performed using the time-dependent model of DEBay. The estimated average NGECs and the actual NGECs for different cell types are shown in (c) and (d), respectively. e) Distribution of the estimated time-dependent NGECs. Time points are represented by increasing order of color intensities. The lowest intensity denotes the initial time point, and the highest intensity denotes the end time point.

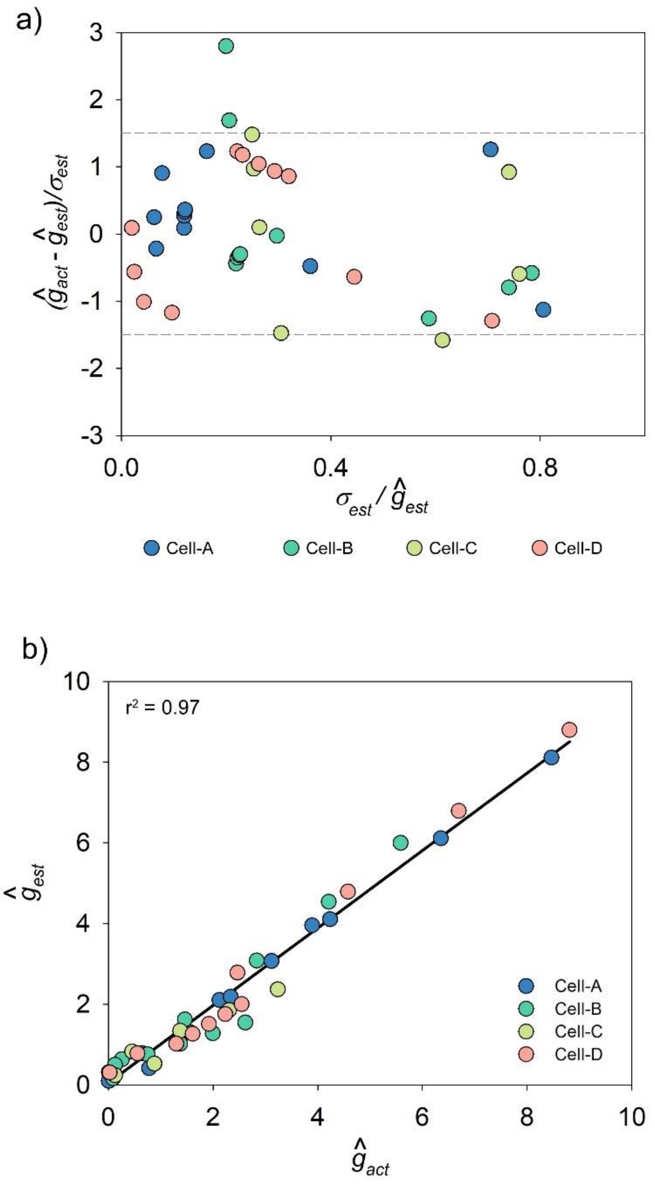


Figure S5: Evaluation of the performance of DEBay for all the time-dependent synthetic data sets. a) The deviation between the mean of estimated NGECs ($\hat{g}_{est}$) and the actual NGECs ($\hat{g}_{act}$) to the standard deviation of the estimated NGECs ($\sigma_{est}$). b) Correlation between the mean of estimated NGECs ($\hat{g}_{est}$) and the actual NGECs ($\hat{g}_{act}$).
